# Changes in subjective preference do not require dopamine signaling or the orbital frontal cortex

**DOI:** 10.1101/865436

**Authors:** Merridee J. Lefner, Alexa P. Magnon, James M. Gutierrez, Matthew R. Lopez, Matthew J. Wanat

**Affiliations:** Neurosciences Institute and Department of Biology, University of Texas at San Antonio, San Antonio, TX 78249, USA

**Author notes:** Corresponding Author: Matthew J. Wanat, Neurosciences Institute, Department of Biology, University of Texas at San Antonio, One UTSA Circle, San Antonio, TX 78249.

## Abstract

‘Sunk’ or unrecoverable costs impact proximal reward-based decisions across species. However, it is not known if these incurred costs elicit a long-lasting change in reward value. To address this, we identified the relative preference between different flavored food pellets in rats. Animals were then trained to experience the initially preferred reward after short delays and the initially less preferred reward after long delays. This training regimen enhanced the preference for the initially less desirable food reward. We probed whether this change in subjective preference involved dopamine signaling or the orbital frontal cortex (OFC) given that these neural systems contribute to reward valuation. Systemic dopamine receptor antagonism attenuated anticipatory responding during training sessions but did not prevent the change in reward preference elicited by incurred temporal costs. OFC lesions had no effect on anticipatory responding during training or on the change in reward preference. These findings collectively illustrate that the neural systems involved with economic assessments of reward value are not contributing to changes in subjective preference.

## Introduction

Reward-based decisions are driven by cost-benefit analyses. A purely economic decision should be influenced solely by the *future* costs associated with earning a reward^1,2^. However, behavioral evidence across species demonstrates that *past* costs influence decision-making policies^3-10^. These ‘sunk costs’ can result in fewer rewards earned, as subjects will persist in an action even though it is advantageous to disengage from their current behavior^9,10^. The effect of sunk costs on behavior has been primarily studied in tasks that involve opportunity costs, in which choosing one course of action comes at the expense of an alternative outcome. As such, it is not known if sunk costs increase the value of the chosen reward or decrease the value of a potential alternative reward. Furthermore, it is not clear if sunk costs only impact proximal decisions or if they elicit long-lasting changes in reward value.

Value-based decisions involve the mesolimbic dopamine system and the orbital frontal cortex (OFC)^11-14^. Dopamine neurons in the ventral midbrain and pyramidal neurons in the OFC encode value-based parameters such as reward size, reward probability, and reward rate^15-19^. Furthermore, economic decisions are altered by antagonizing dopamine receptors in the ventral striatum or lesioning the OFC^20,21^. Therefore, mesolimbic dopamine and the OFC are candidate neural systems that could mediate the influence of sunk costs on behavior.

In this study we examined how sunk costs affect reward preference in male and female rats. To address this, we first identified the relative preference between different flavored food rewards in a free-feeding test. Rats then underwent training sessions in which the initially preferred reward was delivered after short delays (low temporal cost), while the initially less preferred reward was delivered after long delays (high temporal cost). In this manner, we could determine if the high temporal costs that preceded the initially less preferred option subsequently enhanced the preference for that reward. Additionally, we performed pharmacological manipulations and site-specific lesions to establish if the behavioral changes elicited by sunk temporal costs required dopamine signaling or the OFC.

## Methods

### Subjects and surgery

All procedures were approved by the Institutional Animal Care and Use Committee at the University of Texas at San Antonio. Male and female Sprague-Dawley rats (Charles River, MA) were pair-housed upon arrival and given ad libitum access to water and chow and maintained on a 12-hour light/dark cycle. OFC lesion surgeries or sham control surgeries were performed under isoflurane anesthesia on rats weighing between 300-350 g. Rats received injections of NMDA (12.5 µg/µL; Tocris) in saline vehicle at a rate of 0.1 µL/min at the following locations relative to bregma: 3.0 mm anterior, ± 3.2 mm lateral, 5.2 mm ventral (0.05 µL injection); 3.0 mm anterior, ± 4.2 mm lateral, 5.2 mm ventral (0.1 µL injection); 4.0 mm anterior, ± 2.2 mm lateral, 3.8 mm ventral (0.1 µL injection); 4.0 mm anterior, ± 3.7 mm lateral, 3.8 mm ventral (0.1 µL injection)^22^. Sham surgeries involved lowering the injector to the injection site^22^. Animals were allowed to recover for at least 1 week following surgery before beginning training.

### Training

Rats were placed and maintained on mild food restriction (∼15 g/day of standard lab chow) to target 90% free-feeding weight, allowing for an increase of 1.5% per week. Behavioral sessions were performed in chambers that had grid floors, a house light, and two food trays located on a single wall. Only one behavioral session was performed per day. In free-feeding sessions, plastic barriers were placed over the food trays. Additionally, a plastic insert was placed over the grid floors that contained two fixed cups in which the food pellets were placed. Experimental 45-mg sucrose food pellets (chocolate flavor #F0025 and banana flavor #F0024; Bio-Serv) were placed in their home cages to minimize neophobia prior to training. Rats first underwent a free-feeding session (10 mins) in which a single food pellet flavor was offered (6.5 g total). On the following day, rats would undergo a second free-feeding session in which the alternate food pellet flavor was offered (ordering counterbalanced between animals). We would repeat the training on the subsequent day on the rare occasion in which rats did not consume any food during the single reward free-feeding session. Training the rats in this manner ensured animals consumed both of the reward options prior to the preference test.

For the free-feeding preference test, both chocolate and banana food pellets were freely available in cups affixed to the floor. To ensure that there was an ample supply of food, we provided 13 g of each flavor, which was 3 g higher than the maximal amount consumed in pilot studies. We identified which food pellet favor was the Initial Preferred and the Initial Less Preferred reward based on this preference test. Rats next underwent training sessions (1 per day) in which one of the food rewards was delivered non-contingently for a total of 50 pellets per session. The Initial Preferred reward was delivered after a 30 ± 5 s inter-trial interval (ITI) in Short Delay training sessions. The Initial Less Preferred reward was delivered after a 60 ± 5 s ITI in Long Delay training sessions. Rats underwent a total of 10 training sessions, which alternated between Short and Long Delay sessions, with the first session counterbalanced between animals. Rats underwent a second free-feeding preference test following this training regimen (**Fig. 1A**). In a control experiment, rats were trained as described above except that a 45 ± 5 s ITI was utilized for both Initial Preferred and the Initial Less Preferred reward training sessions. For dopamine receptor antagonist experiments, rats received an i.p. injection of flupentixol (225 µg/kg) or saline vehicle 1 hour prior to training sessions.

**Figure 1.**
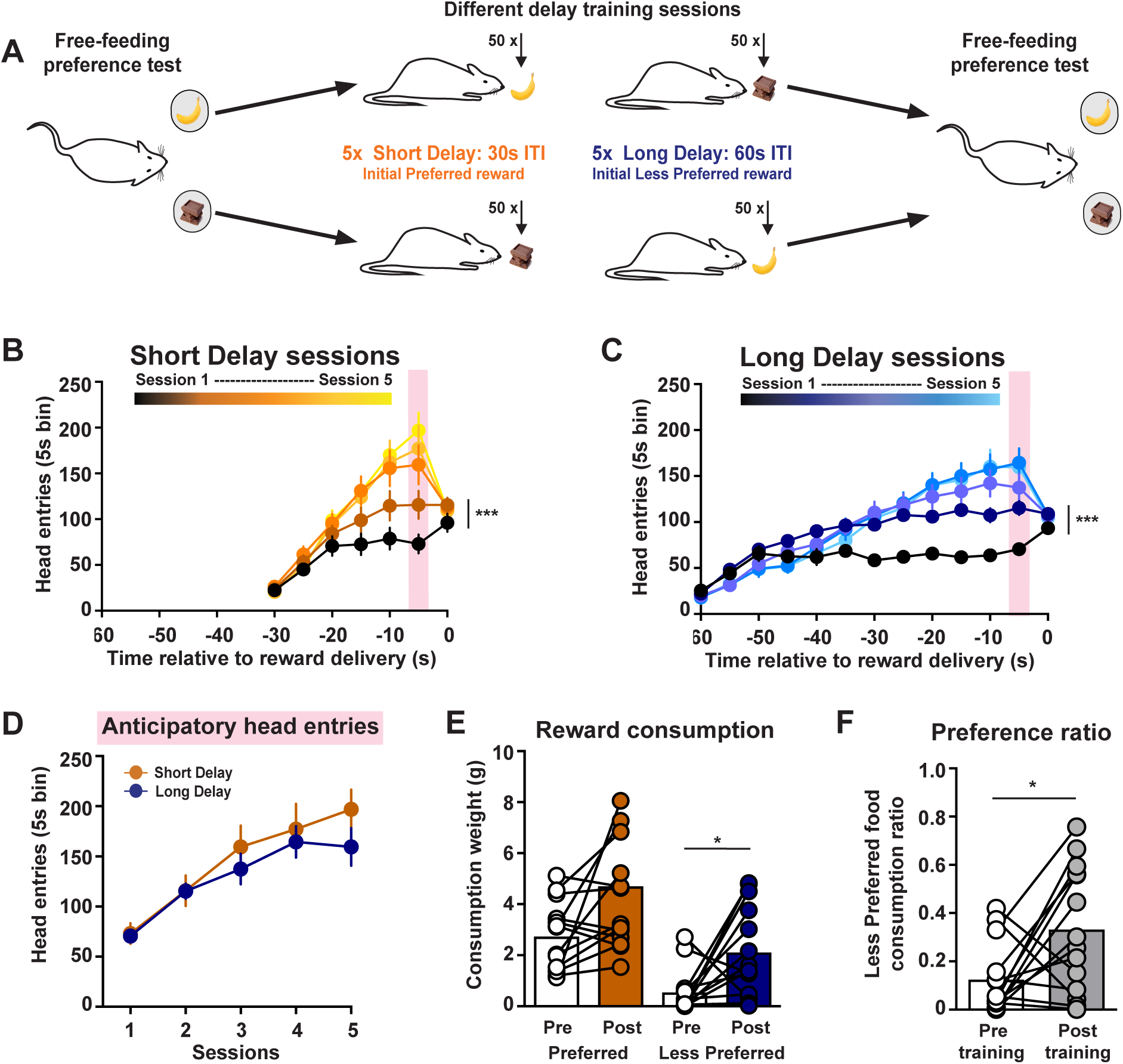
Increased preference for rewards delivered after high temporal costs. (A) Training schematic. (B,C) Head entries into food port across training sessions for Short Delay and Long Delay training sessions. (D) Anticipatory head entries across training sessions. (E) Food consumed during the free-feeding preference tests. (F) Preference ratio. * *p* < 0.05, *** *p* < 0.001.

### Data analysis

Head entries into the food port were quantified in 5 s bins during training sessions. Anticipatory head entries were quantified as the head entries performed in the 5 s preceding the food pellet delivery. The preference ratio was calculated as the amount of the Initial Less Preferred reward consumed relative to the total food consumed during the free-feeding test session. The effect of training on the preference ratio was assessed using a paired t-test. All other statistical analyses utilized a two-way ANOVA (repeated measures where appropriate) followed by a post-hoc Sidak’s multiple comparison test. The Geisser-Greenhouse correction was applied to address unequal variances between groups.

## Results

We hypothesize that the sunk costs which precede a reward influence how the reward is valued. Specifically, we propose that high temporal costs can increase the preference for an intrinsically less desirable reward option. To address this, rats first underwent a 10 min free-feeding preference test between food rewards that had an identical nutritional profile but differed in flavor (chocolate or banana). We identified which was the Initial Preferred and the Initial Less Preferred flavor for each rat based on the food consumed during this free-feeding test. Rats then underwent training sessions in which one of the food rewards was delivered non-contingently for a total of 50 pellets per session. The Initial Preferred reward was delivered after a 30 ± 5 s ITI in Short Delay training sessions. The Initial Less Preferred reward was delivered after a 60 ± 5 s ITI in Long Delay training sessions. Rats underwent a total of 10 training sessions, which alternated between Short and Long Delay sessions, with the first session counterbalanced between animals. Rats then underwent a second free-feeding preference test following this training regimen (**Fig. 1A**).

Food pellets were delivered non-contingently and were not preceded by reward-predictive cues during training sessions. Although no operant action was required, rats increased the number head entries into the food port across Short Delay training (2-way ANOVA: session effect *F*_(4, 52)_ = 6.81, *p* = 0.0002; **Fig. 1B**) and Long Delay training (session effect: *F*_(4, 52)_ = 6.78, *p* = 0.0002; n = 14 rats; **Fig. 1C**). These anticipatory responses peaked prior to the reward delivery and were no different between Short and Long Delay sessions (2-way ANOVA: training effect *F*_(1, 13)_ = 3.28, *p =* 0.093; **Fig. 1D**). This training regimen selectively increased the consumption of the Initial Less Preferred reward (2-way ANOVA: training effect *F*_(1, 13)_ = 25.63, *p* = 0.0002; flavor effect *F*_(1, 13)_ = 17.12, *p* = 0.0012; post-hoc Sidak’s *t*_(13)_ = 2.51, *p* = 0.036; **Fig. 1E**). These findings illustrate that the high temporal costs during training sessions enhanced the value of the Initial Less Preferred reward, without affecting the value of the Initial Preferred reward. As a consequence, rats exhibited an increased preference for the Initial Less Preferred reward following training sessions (paired t-test: *t*_(13)_ = 2.49, *p =* 0.027; **Fig. 1F**).

The change in reward preference could potentially result from the increased exposure to the Initially Less Preferred reward over training sessions. To address this possibility, we trained a separate group of rats as described above except that an identical 45 ± 5 s ITI was used for both the Initial Preferred and Less Preferred reward training sessions (**Fig. 2A**). Rats increased head entries into the food port across sessions for the Initial Preferred reward (2-way ANOVA: session effect *F*_(4, 24)_ = 24.25, *p* < 0.0001; n = 7 rats; **Fig. 2B**) and the Initial Less Preferred reward (session effect *F*_(4, 24)_ = 13.07, *p* < 0.0001; **Fig. 2C**). There was no difference in anticipatory responding between the sessions for the different flavored food rewards (2-way ANOVA: training effect *F*_(1, 6)_ = 1.20, *p =* 0.31; **Fig. 2D**). This training regimen increased the consumption of the Initial Preferred reward in the preference test (2-way ANOVA: training effect *F*_(1, 6)_ = 33.55, *p* = 0.0012; flavor effect *F*_(1, 6)_ = 23.47, *p* = 0.0029; post-hoc Sidak’s *t*_(13)_ = 3.71, *p* = 0.019; **Fig. 2E**). Importantly, there was no change in the relative preference between rewards (paired t-test: *t*_(6)_ = 0.91, *p =* 0.40; **Fig. 2F**). Collectively, these experiments demonstrate that the enhanced preference for the Initial Less Preferred reward was due to long delay training sessions (**Fig. 1**) and not a result of increased experience with that reward over training (**Fig. 2**).

**Figure 2.**
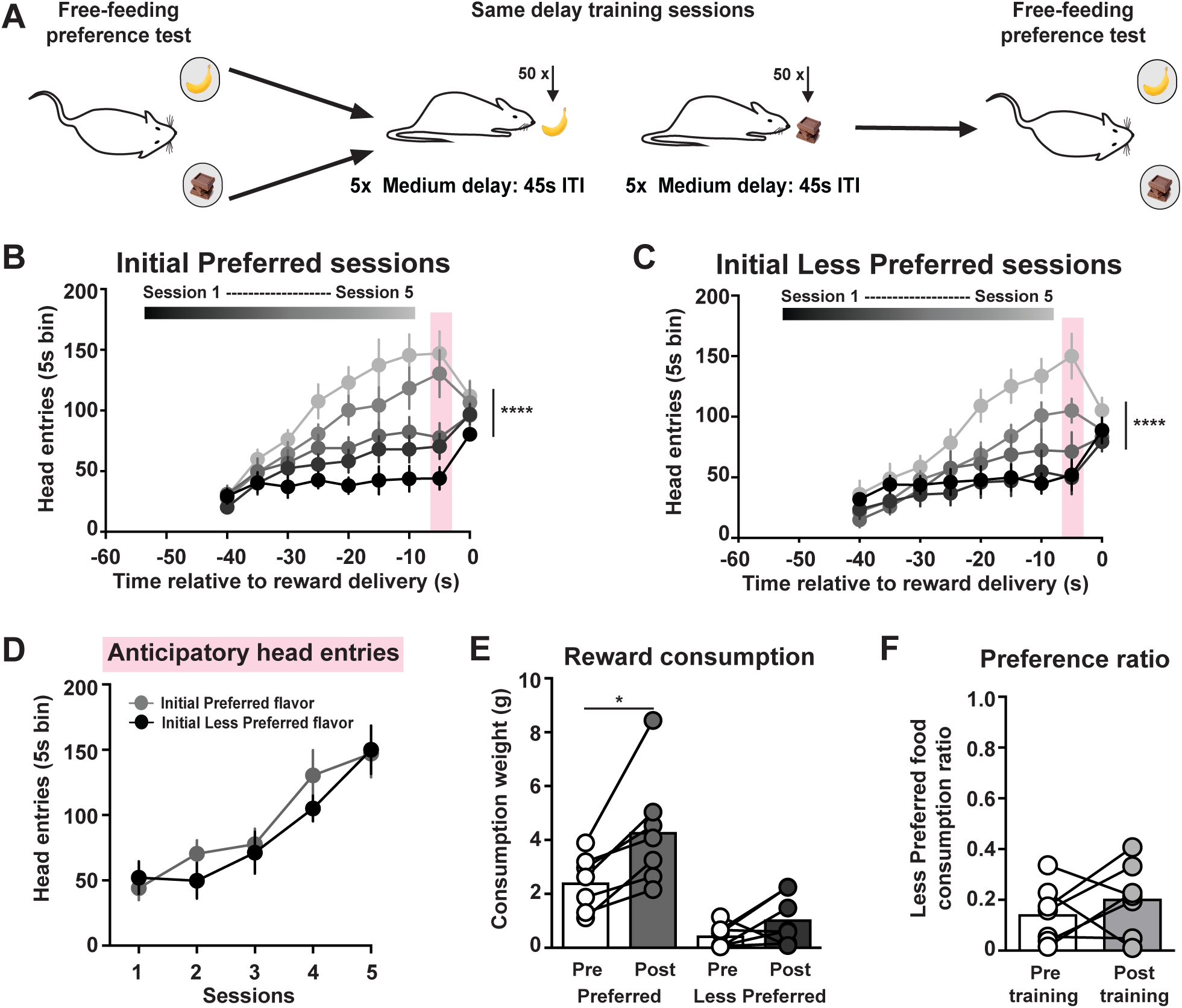
No change in subjective preference following same delay training sessions. (A) Training schematic. (B,C) Head entries into the food port across training sessions for Short Delay and Long Delay training sessions. (D) Anticipatory head entries across training sessions. (E) Food consumed during the free-feeding preference tests. (F) Preference ratio. * *p* < 0.05, **** *p* < 0.0001.

Dopamine neurons respond to rewards to convey the relative preference between options^15,17,23,24^. However, this reward-evoked dopamine response scales with increasing temporal uncertainty^25-27^. Therefore, the increased preference for rewards that follow greater delays could be mediated by dopamine signaling. To address this possibility, we systemically administered flupentixol, a D1/D2 dopamine receptor antagonist, prior to training sessions and examined the consequence on reward preference (**Fig. 3A**). Rats receiving saline injections increased the total number of head entries across Short Delay training (2-way ANOVA: Short Delay session effect *F*_(1, 11)_ = 5.06, *p* = 0.046; Long Delay session effect *F*_(1, 11)_ = 1.59, *p* = 0.23; n = 12 rats; **Fig. 3B**). This increase in head entries was absent in rats that received flupentixol injections (Short Delay session effect *F*_(1, 11)_ = 4.47, *p* = 0.058; Long Delay session effect *F*_(1, 11)_ = 0.9, *p* = 0.36; n = 12 rats; **Fig. 3C**). Furthermore, antagonizing dopamine receptors prevented the emergence of anticipatory responding across both Short Delay sessions (treatment effect *F*_(1, 22)_ = 8.5, *p* = 0.0079; **Fig. 3D**) and Long Delay sessions (treatment effect *F*_(1, 22)_ = 13.26, *p* = 0.0014; **Fig. 3E**).

**Figure 3.**
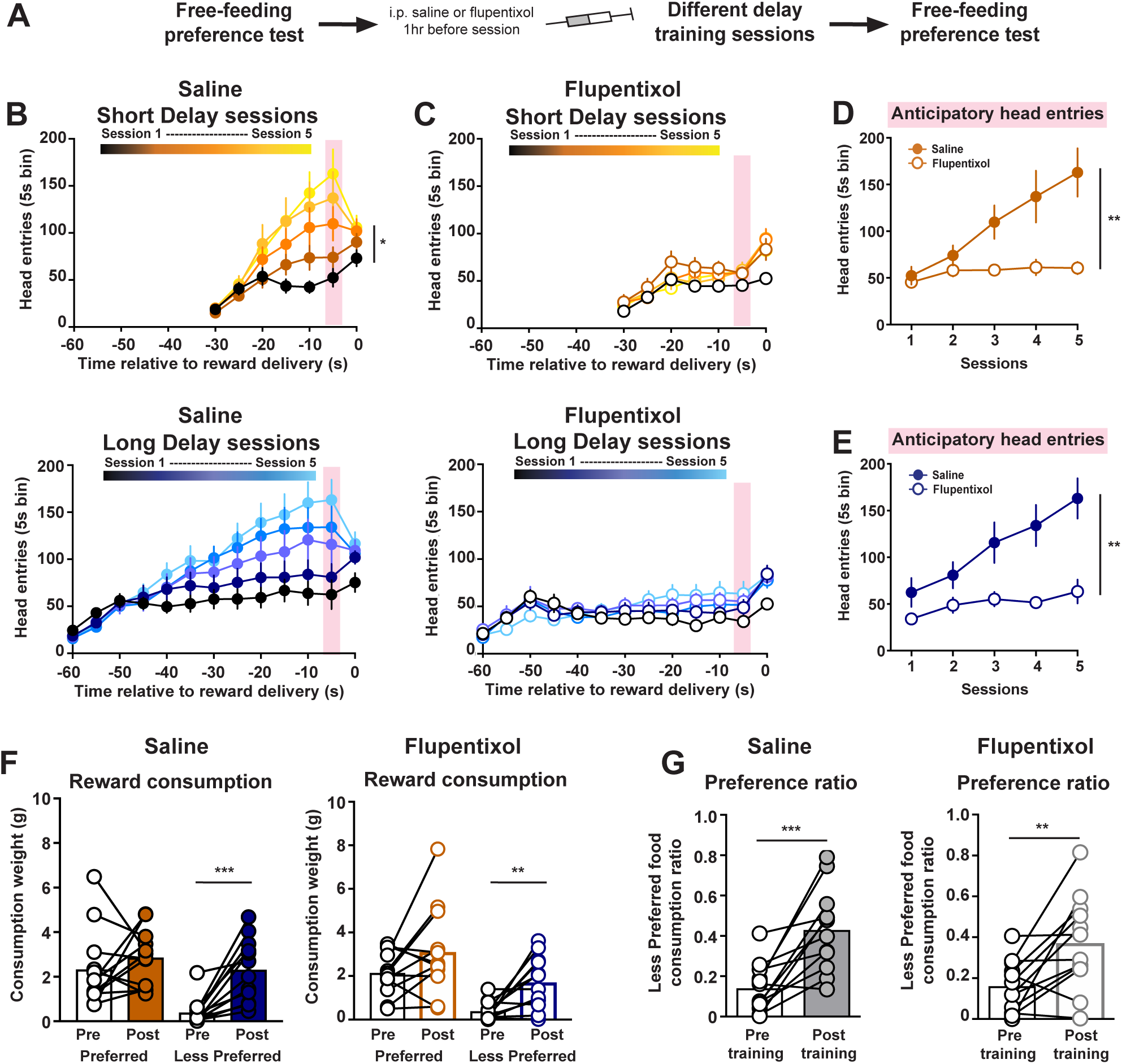
Changes in subjective preference do not involve dopamine signaling. (A) Training schematic. (B) Head entries into food port during Short Delay and Long Delay training sessions in rats receiving saline injections. (C) Head entries into food port during Short Delay and Long Delay training sessions in rats receiving flupentixol injections. (D) Anticipatory responding across Short Delay training sessions in rats receiving saline or flupentixol injections. (E) Anticipatory responding across Long Delay training sessions in rats receiving saline or flupentixol injections. (F) Food consumed during the free-feeding preference tests in saline- and flupentixol-treated rats. (G) Preference ratio in saline- and flupentixol-treated rats. * *p* < 0.05, ** *p* < 0.01, *** *p* < 0.001.

Consistent with our results from untreated animals (**Fig. 1**), saline-treated rats selectively increased the consumption of the Initial Less Preferred reward following training (2-way ANOVA: training effect *F*_(1, 11)_ = 24.16, *p* = 0.0005; flavor effect *F*_(1, 11)_ = 11.75, *p* = 0.0056; post-hoc Sidak’s *t*_(11)_ = 4.92, *p* = 0.0009; **Fig. 3F**). Rats that received flupentixol treatment during training sessions also increased the consumption of the Initial Less Preferred reward (training effect *F*_(1, 11)_ = 17.25, *p* = 0.0016; flavor effect *F*_(1, 11)_ = 12.56, *p* = 0.0046; post-hoc Sidak’s *t*_(11)_ = 3.54, *p* = 0.0093; **Fig. 3F**). Furthermore, both saline- and flupentixol-treated rats exhibited an enhanced preference for the Initial Less Preferred reward (paired t-test: saline *t*_(11)_ = 4.64, *p =* 0.0007; flupentixol *t*_(11)_ = 3.24, *p =* 0.0078; **Fig. 3G**). These data collectively demonstrate that the dopamine system regulates anticipatory responding during training sessions, but does not mediate the change in subjective preference.

The OFC contributes to various aspects of economic decision-making by conveying the value of potential and chosen rewards^12-14,16^. As such the OFC is another candidate region for mediating the change in subjective preference elicited by incurred temporal costs. To address this, we performed excitotoxic lesions of the OFC or sham surgeries prior to the initial preference test (**Fig. 4A**). As expected, sham-treated rats increased head entries across Short Delay training (2-way ANOVA: session effect *F*_(4, 36)_ = 3.92, *p* = 0.0096) and Long Delay training (2-way ANOVA: session effect *F*_(4, 36)_ = 3.57, *p* = 0.015; n = 10 rats; **Fig. 4B**). Rats with OFC lesions also increased the total number of head entries across training sessions (Short Delay: session effect *F*_(4, 36)_ = 3.34, *p* = 0.02; Long Delay: session effect *F*_(4, 36)_ = 3.98, *p* = 0.009; **Fig. 4C**). There was no difference in anticipatory responding between sham and OFC-lesioned rats during Short Delay sessions (treatment effect *F*_(1, 18)_ = 1.42, *p* = 0.25; **Fig. 4D**) and Long Delay sessions (treatment effect *F*_(1, 18)_ = 1.17, *p* = 0.29; **Fig. 4E**).

**Figure 4.**
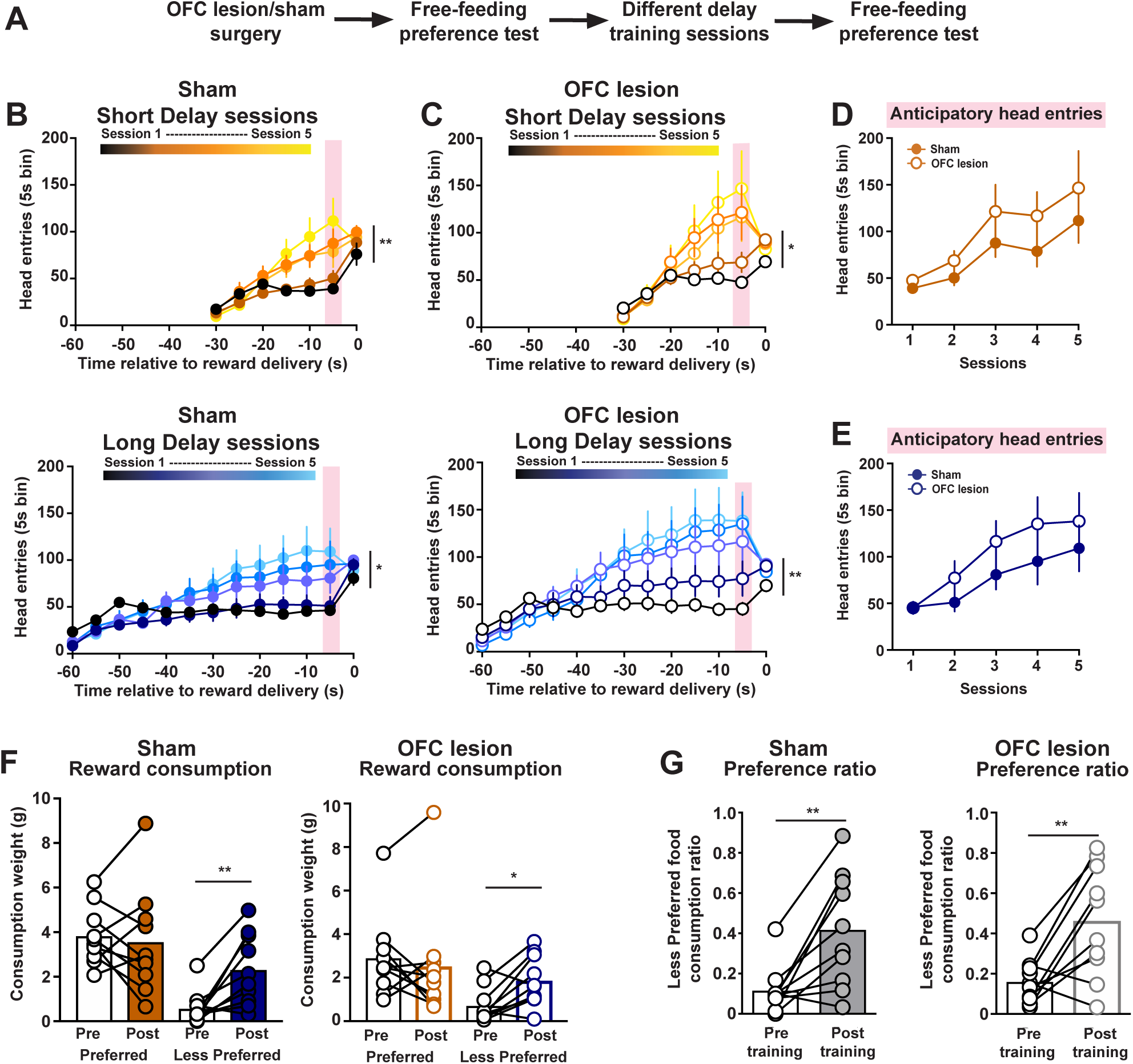
The OFC is not required for changes in subjective preference. (A) Training schematic. B) Head entries into food port during Short Delay and Long Delay training sessions in sham rats. (C) Head entries into food port during Short Delay and Long Delay training sessions in OFC lesioned rats. (D) Anticipatory responding across Short Delay training sessions in sham and OFC lesioned rats. (E) Anticipatory responding across Long Delay training sessions in sham and OFC lesioned rats. (F) Food consumed during the free-feeding preference tests in sham and OFC lesioned rats. (G) Preference ratio in sham and OFC lesioned rats. * *p* < 0.05, ** *p* < 0.01.

We next examined whether OFC lesions altered the change in food consumption and reward preference following training. Sham-treated rats increased the consumption of the Initial Less Preferred reward (2-way ANOVA: training effect *F*_(1, 9)_ = 14.88, *p* = 0.0039; flavor effect *F*_(1, 9)_ = 8.608, *p* = 0.0166; post-hoc Sidak’s *t*_(9)_ = 3.87, *p* = 0.0074; **Fig. 4F**). Similarly, we found that OFC-lesions rats also selectively increased the consumption of the Initial Less Preferred reward (2-way ANOVA: training effect *F*_(1, 9)_ = 3.347, *p* = 0.1006; flavor effect *F*_(1, 9)_ = 4.727, *p* = 0.0577; post-hoc Sidak’s *t*_(9)_ = 3.26, *p* = 0.0195; **Fig. 4F**). Furthermore, both sham-treated and OFC-lesioned rats exhibited an enhanced preference for the Initial Less Preferred reward following training sessions (paired t-test: sham *t*_(9)_ = 3.894, *p =* 0.0037; OFC lesion *t*_(9)_ = 3.685, *p =* 0.005; **Fig. 4G**). Collectively, these results illustrate that neural systems involved with economic decision-making processes, such as the mesocorticolimbic dopamine system and the OFC, are not contributing to changes in subjective preference.

## Discussion

Future actions are guided by one’s past experience. Behavioral evidence in rodents, pigeons, and humans demonstrate that past costs influence reward-based decisions^3-10^. These sunk costs can result in subjects persisting in a suboptimal action, even though it would be advantageous to disengage from their current behavior^9,10^. The impact of sunk costs is commonly assessed using decision making tasks, which cannot determine if past costs increase the value of the chosen option or decrease the value of the alternative option. Here, we developed a training procedure that assessed how past temporal costs influence reward value in a free-feeding preference assay. Our results demonstrate that high temporal costs that precede a reward can subsequently increase the preference for that reward. Therefore, the impact of past costs is not just limited to proximal decisions, as we find that incurred temporal costs can subsequently alter reward preference in a different context.

The relative preference between reward options is linked to the difference in how rewards are valued. This assessment of reward value depends on subjective factors such as one’s internal state (e.g. satiety) and economic parameters, such as reward size^11,28^. Based on the food consumed during the free-feeding preference test, we could identify the relative value between different flavored food rewards as well as changes in reward value following training. Rats that were trained to experience the Initial Less Preferred reward following long delays and the Initial Preferred reward following short delays selectively increased the consumption of the Initial Less Preferred option. Prior research has identified a number of factors that can alter reward preference, including changing the reward size and inducing state-specific satiety^23,29-31^. Our current findings have uncovered that past temporal costs are an additional factor that influences reward preference.

Dopamine neuron firing to the delivery of rewards conveys subjective preference^24^. Furthermore, this reward-evoked dopamine response updates with changes in reward value^15,23^, which together indicates dopamine signaling could contribute to changes in subjective preference. However, our data argues against this possibility, as systemically antagonizing dopamine receptors did not prevent the change in reward preference elicited by incurred temporal costs. Despite the lack of an effect on subjective preference, dopamine signaling is required for the emergence of anticipatory responding.

The OFC contributes to reinforcer valuation^12,13,16^, and is therefore also well-positioned to mediate changes in subjective preference. However, OFC lesions did not alter anticipatory responding during training sessions or prevent the increased preference for the initial less preferred reward. We note that future studies are needed to identify the brain regions responsible for the change in subjective preference elicited by incurred temporal costs. Regardless, our current findings along with prior research indicates that value-based economic decisions and changes in subjective preference are mediated by distinct neural systems.

## Acknowledgements

This work was supported by National Institutes of Health grants DA033386 (MJW) and DA042362 (MJW). The authors declare no conflict of interest.

## Author Contributions

MJL, APM, JMG, MRL performed the experiments and analyzed the data. MJL and MJW designed the experiments and wrote the manuscript.

